# Xenon and nitrous oxide induced changes in resting EEG activity can be explained by systematic increases in the relaxation rates of stochastically driven alpha band oscillatory activity

**DOI:** 10.1101/2024.07.07.602148

**Authors:** Rick Evertz, Andria Pelentritou, John Cormack, Damien G. Hicks, David T. J. Liley

## Abstract

Resting electroencephalographic activity is typically indistinguishable from a filtered linear random process across a diverse range of behavioural and pharmacological states, suggesting that the power spectral density of the resting electroencephalogram (EEG) can be modelled as the superposition of multiple, stochastically driven and independent, alpha band (8 – 13 Hz) relaxation oscillators. This simple model can account for variations in alpha band power and ‘1/f scaling’ in eyes-open/eyes-closed conditions in terms of alterations in the distribution of the alpha band oscillatory relaxation rates. As changes in alpha band power and ‘1/f scaling’ have been reported in anaesthesia we hypothesise that such changes may also be accounted for by alterations in alpha band relaxation oscillatory rate distributions. On this basis we choose to study the EEG activity of xenon and nitrous oxide, gaseous anaesthetic agents that have been reported to produce different EEG effects, notable given they are both regarded as principally acting via N-methyl-D-aspartate (NMDA) receptor antagonism. By recording high density EEG from participants receiving equilibrated step-level increases in inhaled concentrations of xenon (n = 24) and nitrous oxide (n = 20), alpha band relaxation rate (damping) distributions were estimated by solving an inhomogeneous integral equation describing the linear superposition of multiple alpha-band relaxation oscillators having a continuous distribution of dampings. For both agents, level-dependent reductions in alpha band power and spectral slope exponent (15-40 Hz) were observed, that were accountable by increases in mean alpha band damping. These shared increases suggest that, consistent with their identified molecular targets of action, xenon and nitrous oxide are mechanistically similar, a conclusion further supported by neuronal population modelling in which NMDA antagonism is associated with increases in damping and reductions in peak alpha frequency. Alpha band damping may provide an important link between experiment and theories of consciousness, such as the global neuronal network theory, where the likelihood of a globally excited state (‘conscious percept’), is inversely related to mean damping.

## Introduction

The gases xenon and nitrous oxide are able to induce dissociative effects at sub-anaesthetic levels and are both ostensibly N-methyl-D-aspartate (NMDA) receptor antagonists. However, they each produce a different constellation of effects on the magnetoencephalogram (MEG) and electroencephalogram (EEG). For example, xenon while typically reported to induce increases in low-frequency delta (0 – 4 Hz) and theta (4 – 8 Hz) band activity, is associated with inconsistent changes in higher frequency alpha (8 – 13 Hz), beta (13 – 30 Hz) and gamma activity (*>* 30 Hz) [1–6]. In contrast nitrous oxide is reported to have little or no effect on low frequency delta and theta activity, but is generally identified as suppressing alpha and increasing high frequency gamma [2, 7–12].

Pelentritou et al [2] concluded, based on the most detailed EEG and MEG source level spectral analysis of dose dependent xenon and nitrous oxide effect to date, that “… we find no clear evidence that xenon and nitrous oxide gaseous anesthesia are acting via similar mechanisms, at least when evaluated in terms of dose– and modality-dependent variations in [band defined] spectral power.”

However, as we will demonstrate both xenon and nitrous oxide evince similar dose dependent changes in the power-law spectral scaling of electroencephalographic power between 15 and 40 Hz. Because these shared changes in spectral scaling can be theoretically accounted for in terms of systematic alterations in the damping of stochastically perturbed alpha-band relaxation oscillatory activity, a shared mechanism of action *is* suggested.

Power-law spectral scaling of the EEG is typically thought to indicate the presence of arrhythmic or aperiodic activity that signifies the extent to which the brain is dynamically operating near some critical boundary optimal for information processing [13, 14]. It is considered both dynamically and functionally distinct from the corresponding rhythmic activity (the well defined peaks in the power spectrum), as exemplified by alpha band activity [3, 15–17].

However, an alternative perspective exists, that accords with the empirically established stochastic linear nature of spontaneous EEG and well-established mean field models of EEG electro-rhythmogenesis. In this approach resting state EEG is theorised to arise from the summed activity of cortical populations of stochastically driven linear relaxation oscillators that oscillate about a narrow range of alpha band frequencies [18]. By assuming a distribution of population relaxation rates (dampings) ‘1/F’ spectral scaling emerges in the context of rhythmic alpha oscillatory activity. From this viewpoint nominal rhythmic and arrhythmic activity are no longer functionally and dynamically distinct.

By mathematically formulating the distribution of population relaxation rates as the solution to a Fredholm first order integral equation it has been shown that changes in alpha band power between eyes-closed and eyes-open are the result of an increase in mean alpha band damping, a result consistent with that obtained using fixed-order autoregressive moving average (ARMA) time series modelling [19].

We apply this alternative perspective to the analysis of high-density, sensor level, EEG data recorded during stepwise increases in steady state xenon and nitrous oxide levels in healthy participants. We find that the level induced reductions in alpha band spectral power and flattening of the ‘1/f’ spectral scaling can be well accounted for in terms of systematic changes in the distribution of estimated relaxation rates. Further, by interpreting the dose dependent changes in the estimated damping distributions in terms of the parametric behaviour of a well-known mean field theory of electrocortical activity [20, 21], a mechanism of action consistent with the reported glutamatergic pharmacological site of xenon and nitrous oxide action emerges.

## Results

We detail the quantified, level-dependent, power spectral changes induced by xenon and nitrous oxide, followed by modelling which aims to account for these changes in terms of a sum of stochastically driven alpha band relaxation oscillatory processes having a distribution of relaxation rates (dampings). Finally, we identify shared, level-dependent, changes in the damping distributions.

### Xenon and nitrous oxide level dependent EEG spectral changes

Fig 1a presents normalised histograms of the alpha band (8 – 13 Hz) power and spectral scaling exponent, across all electrodes (64) and participants (24), for the eyes-closed baseline (0%) and xenon end-tidal concentrations of 8%, 16%, 24% and 42%. Significant xenon level-dependent decreases in median absolute alpha band power are observed (xenon alpha band power population median: 18.503 [0%], 16.361 [8%], 15.385 [16%], 14.947 [24%] and 13.384 [42%] *µV* ^2^). Table 1a details pair-wise level differences together with their statistical significance. As expected, the largest absolute change in population median alpha band power was between rest and 42%.

**Fig 1.**
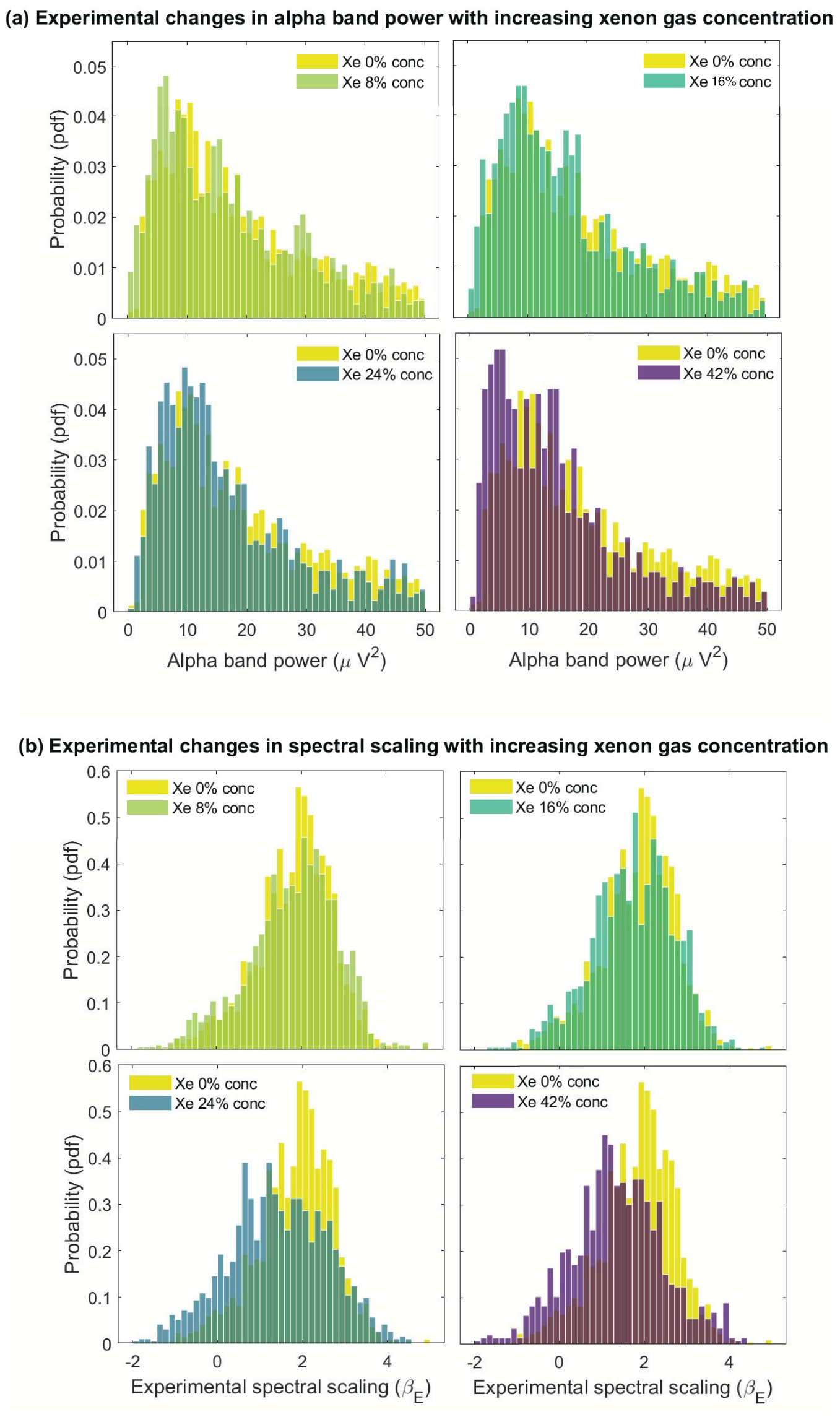
Experimental changes in alpha band power and the spectral scaling exponent (*β*) with increasing levels of xenon. (**a**) Normalised histogram of alpha band (8 – 13 Hz) power, across all electrodes (64) and participants (24), for an eyes-closed baseline and increasing end-tidal concentrations of xenon (0% (baseline), 8%, 16%, 24% and 42%). Minimal changes in the alpha band power with increasing xenon gas concentrations are observed. **(b)** Normalised histogram of the 15 – 40 Hz spectral scaling exponent (*β*), across all electrodes and participants, for increasing levels of xenon gas (0% (baseline), 8%, 16%, 24% and 42%). In contrast to the alpha band power, prominent level dependent changes are observed in the distribution of the spectral scaling exponent.

**Table 1.**
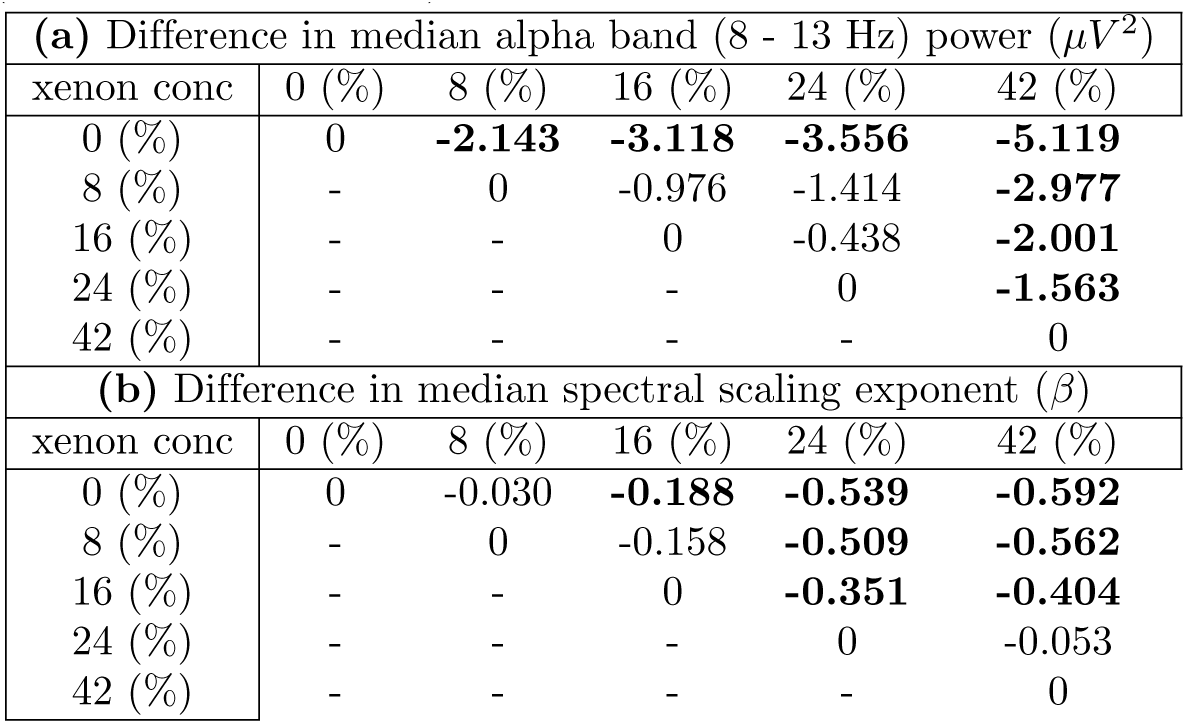
Changes in median alpha band (8 – 13 Hz) power and spectral scaling exponent with increasing levels of xenon. (**a**) Pairwise differences (column – row) in the median value of the distribution of estimated alpha band power (*µV* ^2^) between all conditions (0% (baseline), 8%, 16%, 24% and 42%). **(b)** As for (a) except for pairwise differences calculated across spectral scaling exponent (*β*). Bolded numbers indicate pair-wise statistical significance (*p <* 0.05) corrected for multiple comparisons (see Methods for details).

| **(a) Difference in median alpha band (8 - 13 Hz) power ($\mu V^2$)** |  |  |  |  |  |
| --- | --- | --- | --- | --- | --- |
| xenon conc | 0 (%) | 8 (%) | 16 (%) | 24 (%) | 42 (%) |
| 0 (%) | 0 | **-2.143** | **-3.118** | **-3.556** | **-5.119** |
| 8 (%) | - | 0 | -0.976 | -1.414 | **-2.977** |
| 16 (%) | - | - | 0 | -0.438 | **-2.001** |
| 24 (%) | - | - | - | 0 | **-1.563** |
| 42 (%) | - | - | - | - | 0 |
| **(b) Difference in median spectral scaling exponent ($\beta$)** |  |  |  |  |  |
| xenon conc | 0 (%) | 8 (%) | 16 (%) | 24 (%) | 42 (%) |
| 0 (%) | 0 | -0.030 | **-0.188** | **-0.539** | **-0.592** |
| 8 (%) | - | 0 | -0.158 | **-0.509** | **-0.562** |
| 16 (%) | - | - | 0 | **-0.351** | **-0.404** |
| 24 (%) | - | - | - | 0 | -0.053 |
| 42 (%) | - | - | - | - | 0 |

The spectral scaling slope exponent, *β*, was estimated by fitting a 1*/f^β^* profile to the high frequency tail of the power spectrum (15-40 Hz). A broad range of spectral scaling slope exponents were observed (see Fig 1b) across both baseline and xenon conditions (*−*2 *≤ β ≤* 5). Negative spectral scaling slope exponents (*−*2 *≤ β ≤* 0) occurred due to a range of EEG spectra having more power in higher frequencies than in lower, resulting in a positive slope of the overall power spectral density. Significant xenon level-dependent decreases in the population median spectral scaling exponent were observed (xenon spectral scaling exponent population median: 1.956 [0%], 1.9257 [8%], 1.768 [16%], 1.417 [24%] and 1.364 [42%]). Spectral scaling became more shallow as the level of xenon increased, as indicated by the graded changes in the distribution shape and significant pair-wise changes in the population median (Table 1b).

Fig 2a illustrates normalised histograms for the alpha band (8 – 13 Hz) power and spectral scaling exponent, across all electrodes (64) and participants (20), for the eyes-closed baseline (0%) and end-tidal nitrous oxide concentrations of 16%, 32% and 47%. Similar to xenon, nitrous oxide induced level-dependent changes in the distribution of alpha band power (nitrous oxide alpha band power population median: 21.654 [0%], 15.413 [16%], 13.905 [32%] and 15.588 [47%] *µV* ^2^). As with xenon, nitrous oxide induced statistically significant reductions in total alpha band power between rest and increasing gas concentrations (Table 2a). However, higher levels of gas concentration did not necessarily result in statistically significant stepped reductions in alpha band power as indicated by the pair-wise comparisons of each nitrous oxide level. An end-tidal gas concentration of 32% produced the greatest absolute change in alpha band power from baseline.

**Fig 2.**
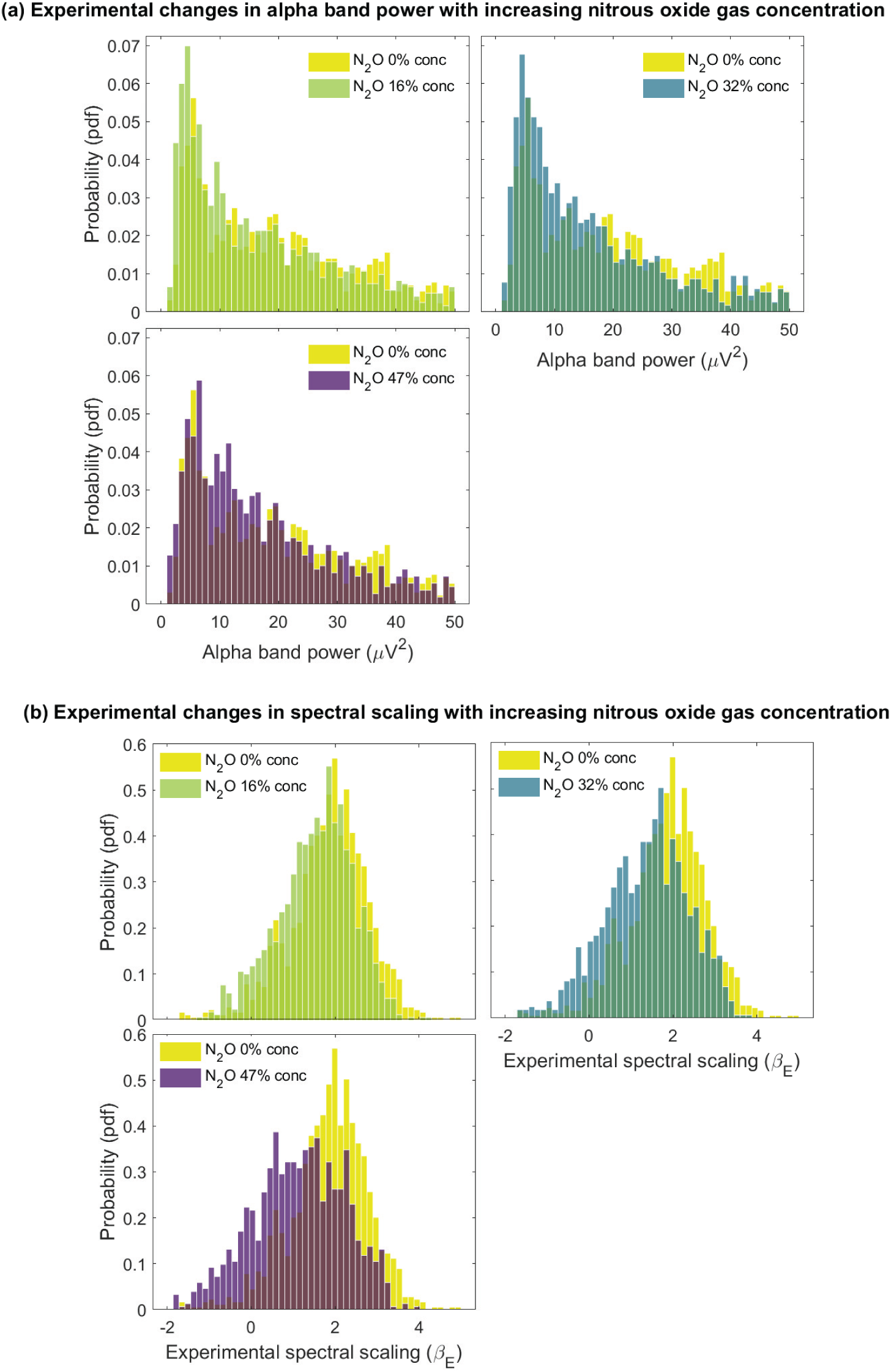
Experimental changes in alpha band power and the spectral scaling exponent (*β*) with increasing levels of nitrous oxide. (**a**) Normalised histogram of alpha band (8 – 13 Hz) power, across all electrodes (64) and participants (20), for an eyes-closed baseline and increasing end-tidal concentrations of nitrous oxide (0% (baseline), 16%, 32% and 47%). Similar to xenon, minimal level dependent changes in the alpha band power are observed. **(b)** Normalised histogram of the 15 – 40 Hz spectral scaling exponent (*β*), across all electrodes and participants, for increasing levels of nitrous oxide gas (0% (baseline), 16%, 32% and 47%). In contrast to alpha band power, and similar to xenon, prominent level dependent changes are observed in the distribution of the spectral scaling exponent.

**Table 2.** Changes in median alpha band power and spectral scaling exponent with increasing levels of nitrous oxide. (**a**) Pairwise differences (column – row) in the median value of the distribution of estimated alpha band (8 – 13 Hz) power (*µV* ^2^) between all conditions (0% (baseline), 16%, 32% and 47%). **(b)** As for (a) except for pairwise differences calculated across spectral scaling exponent (*β*). Bolded numbers indicate pair-wise statistical significance (*p <* 0.05) corrected for multiple comparisons (see Methods for details).

| **(a) Difference in median alpha band (8 -13 Hz) power ($\mu V^2$)** |  |  |  |  |
| --- | --- | --- | --- | --- |
| n2o conc | 0 (%) | 16 (%) | 32 (%) | 47 (%) |
| 0 (%) | 0 | **-6.241** | **-7.749** | **-6.066** |
| 16 (%) | - | 0 | -1.508 | -0.175 |
| 32 (%) | - | - | 0 | -1.683 |
| 47 (%) | - | - | - | 0 |
| **(b) Difference in median spectral scaling exponent ($\beta$)** |  |  |  |  |
| n2o conc | 0 (%) | 16 (%) | 32 (%) | 47 (%) |
| 0 (%) | 0 | **-0.292** | **-0.471** | **-0.762** |
| 16 (%) | - | 0 | **-0.178** | **-0.469** |
| 32 (%) | - | - | 0 | **-0.291** |
| 47 (%) | - | - | - | 0 |

As for xenon, the distribution of spectral scaling exponent (*β*) across the baseline recordings and increasing levels of nitrous oxide displayed a broad range of scaling behaviour (*−*2 *≤ β ≤* 5). Significant reductions in the spectral slope exponent were observed with increasing gas concentration (Fig 2b) (nitrous oxide spectral scaling population median: 1.929 [0%], 1.636 [16%], 1.458 [32%] and 1.167 [42%]). Pair-wise comparisons between rest and increasing gas concentrations revealed that nitrous oxide induces graded reductions in the slope of the power spectrum, with higher gas concentrations resulting in more shallower spectral scaling (Table 2b), a feature consistent with xenon.

Level dependent reductions in peak alpha frequency were observed for both xenon and nitrous oxide. Fig 3a presents the normalised histograms of peak alpha frequency, across all electrodes (64) and participants (24) for eyes-closed baseline (0%) and end-tidal xenon concentrations of 8% 16%, 24% and 42%, with level dependent reductions in median peak alpha frequency observed (xenon peak alpha frequency population median 10.106 [0%], 9.870 [8%], 9.847 [16%], 9.815 [24%] and 9.322 [42%] Hz). Fig 3b presents the corresponding normalised histograms, across all electrodes (64) and participants (20), for nitrous oxide, with a similar sequence of level dependent reductions in median peak alpha frequency observed (nitrous oxide peak alpha frequency population median 10.320 [0%], 10.262 [16%], 10.212 [32%] and 9.978 [47%] Hz). The obvious peaks at 8 Hz arise as a consequence of the lower limit in estimating the peak alpha frequency from fitting a Lorentzian-like profile to the experimental power spectrum (see Methods for further details). Pairwise differences in the median peak alpha frequency and their statistical significance, for both xenon and nitrous oxide, are presented in Tables 3a & b.

**Fig 3.**
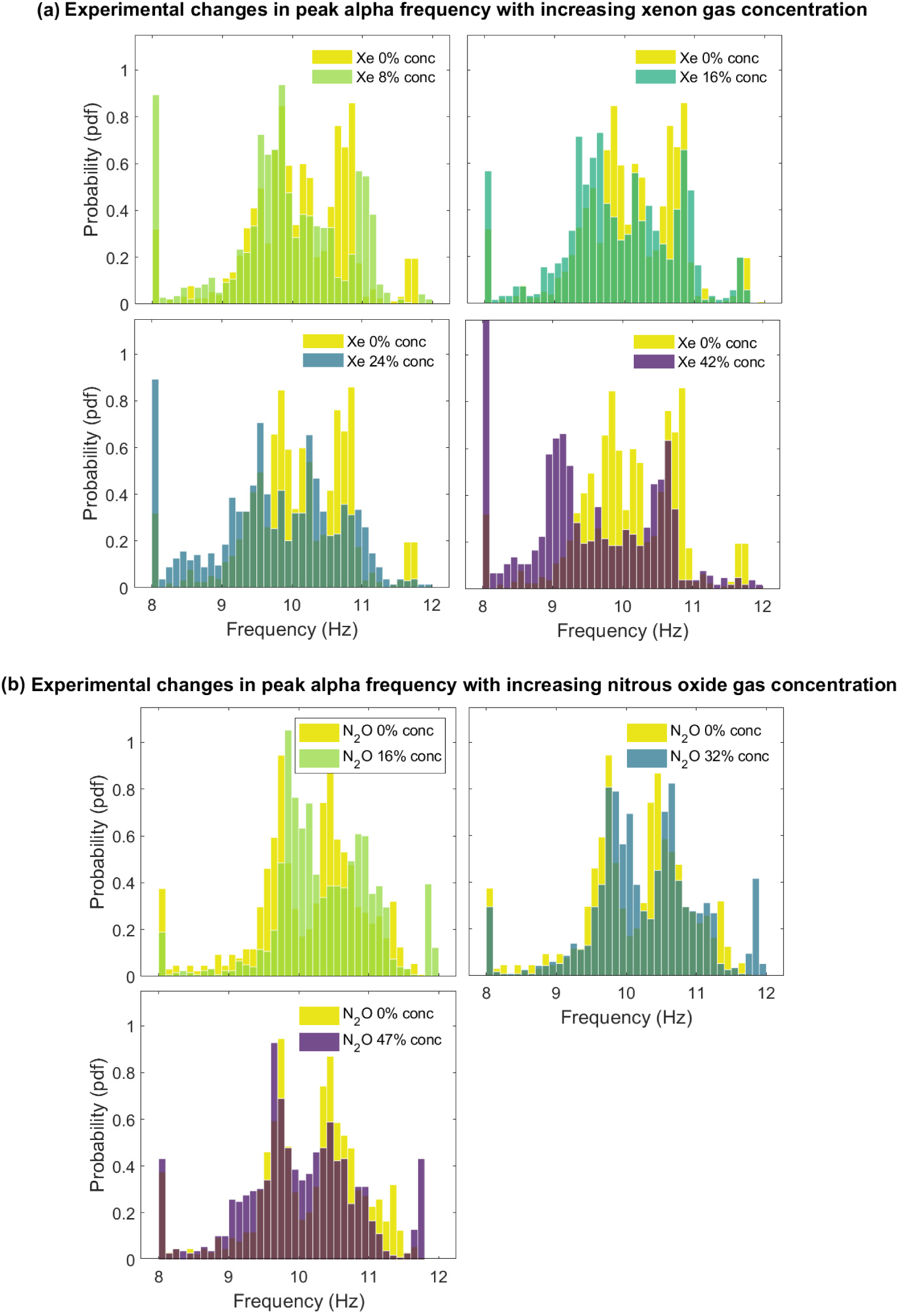
Experimental changes in peak alpha frequency with xenon and nitrous oxide. (**a**) Normalised histogram of peak alpha frequency, across all electrodes (64) and participants (24), for baseline and increasing end-tidal concentrations of xenon gas (0% (baseline), 8%, 16%, 24% and 42%). A clear reduction in peak alpha frequency at the highest xenon gas concentrations is observed. **(b)** Normalised histogram of peak alpha frequency, across all electrodes (64) and participants (20), for baseline and increasing levels of nitrous oxide (0% (baseline), 16%, 32% and 47%). Similar to xenon, reductions in peak alpha frequency are observed at the highest nitrous oxide end-tidal gas concentrations.

**Table 3.** Pairwise differences in the median peak alpha frequency for increasing levels of xenon and nitrous oxide. (**a**) Pairwise differences (column – row) in the population median value for the peak alpha frequency between all xenon conditions (0% (baseline), 8% 16%, 24% and 42%). **(b)** As for (a) except for increasing levels of nitrous oxide (0% (baseline), 16%, 32% and 47%). Bolded numbers indicate pair-wise statistical significance (*p <* 0.05) corrected for multiple comparisons (see Methods for details).

| (a) Differences in xenon median peak alpha frequency (Hz) |  |  |  |  |  |
| --- | --- | --- | --- | --- | --- |
| gas conc | 0 (%) | 8 (%) | 16 (%) | 24 (%) | 42 (%) |
| 0 (%) | 0 | **-0.2360** | **-0.2590** | **-0.2910** | **-0.7810** |
| 8 (%) | - | 0 | -0.0230 | **-0.0550** | **-0.5480** |
| 16 (%) | - | - | 0 | -0.0320 | **-0.5250** |
| 24 (%) | - | - | - | 0 | **-0.4930** |
| 42 (%) | - | - | - | - | 0 |
| (b) Differences in nitrous oxide median peak alpha frequency (Hz) |  |  |  |  |  |
| gas conc | 0 (%) | 16 (%) | 32 (%) | 47 (%) |  |
| 0 (%) | 0 | **-0.0570** | **-0.1080** | **-0.3420** |  |
| 16 (%) | - | 0 | **-0.050** | **-0.2840** |  |
| 32 (%) | - | - | 0 | **-0.2340** |  |
| 47 (%) | - | - | - | 0 |  |

Changes in alpha band power as a function of gas end-tidal concentration were further explored topographically by averaging across participants. Figures 4a & b present the topographic variation of the differences in average alpha band power from baseline (eyes-closed) for xenon and nitrous oxide for increasing end-tidal gas concentration. The pattern of the topographic changes between baseline and each gas level are consistent across both anaesthetic agents and show an occipital/parietal dominance, as expected given the occipital dominance of eyes-closed resting alpha. Similarly, Figures 5a & b illustrate the topographic changes in the spectral scaling exponent. Increases in xenon and nitrous oxide end-tidal gas concentrations elicited, statistically significant and topographically widely distributed, reductions in the spectral scaling exponent. Notably such reductions were more prominent frontally for xenon than for nitrous oxide.

**Fig 4.**
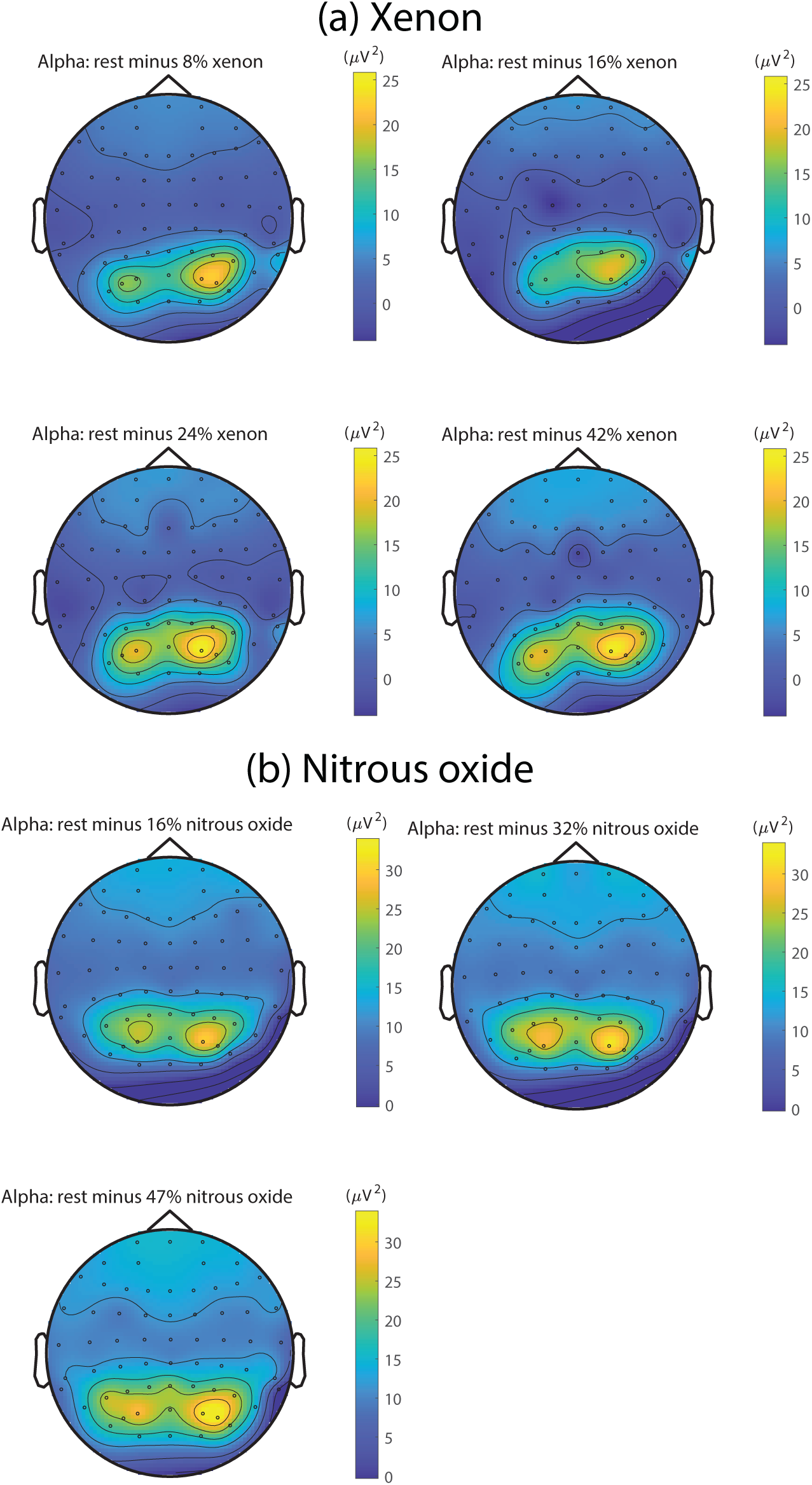
Topographic changes in experimental alpha band power for different levels of xenon and nitrous oxide. (**a**) Average topographic plots, across all participants (24), of the level dependent changes in the alpha band (8 – 13 Hz) power between an eyes-closed baseline and stepwise increasing end-tidal xenon gas concentrations (8%, 16%, 24% and 42%). **(b)** Average topographic plots, across all participants (20), of the level-dependent changes in the alpha band (8 – 13 Hz) power between an eyes-closed baseline and stepwise increasing end-tidal nitrous oxide gas concentrations (16%, 32% and 47%). While xenon and nitrous oxide produce similar averaged changes in the topographic distribution alpha band power none of the level-dependent changes at the group level, for either gas, was statistically significant (as determined via a, false discovery rate corrected, nonparametric permutation test – see Methods for further details).

**Fig 5.**
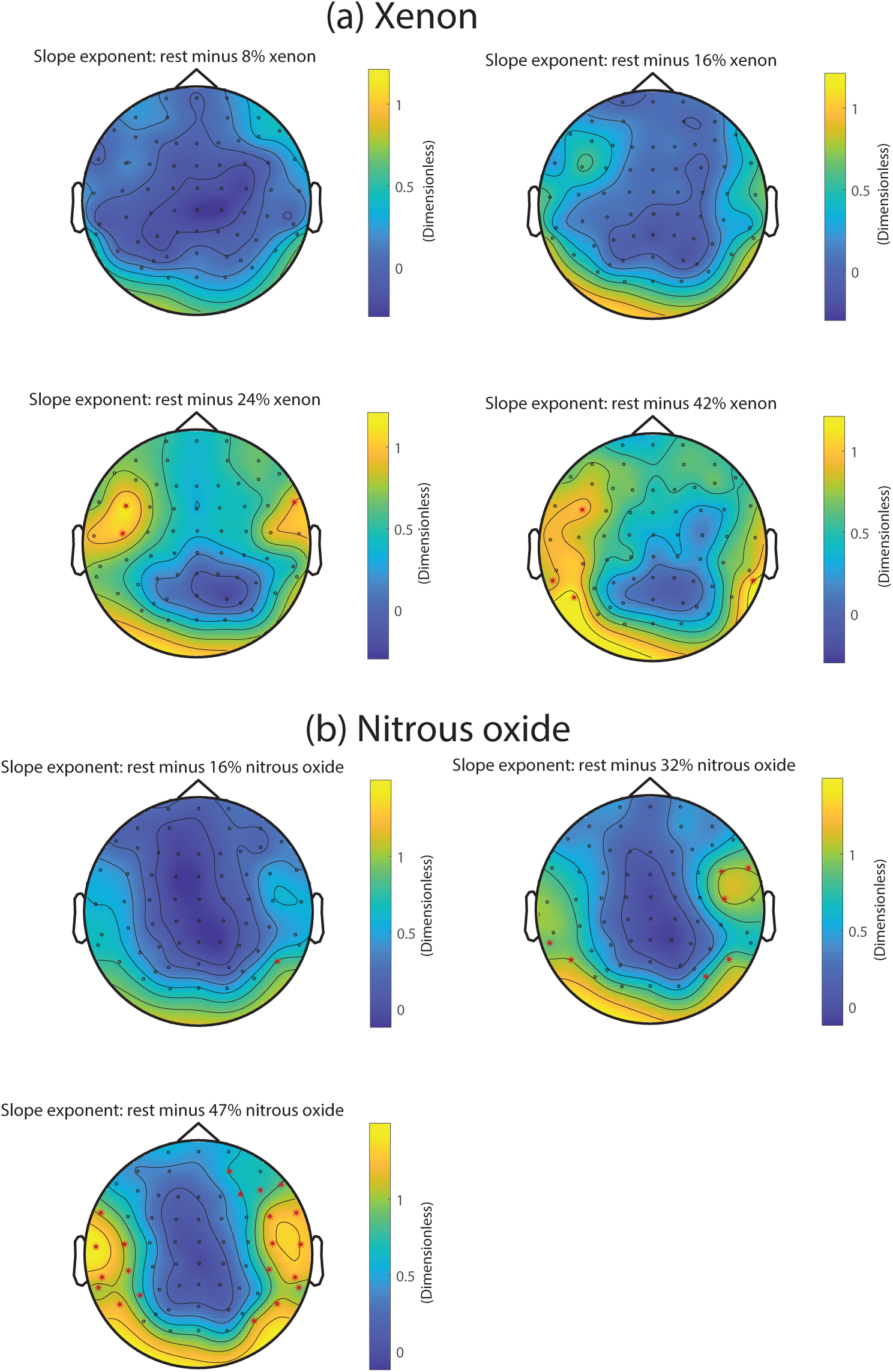
Topographic changes in the experimental spectral scaling exponent (*β*) for different levels of xenon and nitrous oxide. (**a**) Average topographic plots, across all participants (24), of the level dependent changes in the 15 – 40 Hz spectral scaling exponent (*β*) between an eyes-closed baseline and stepwise increasing end-tidal xenon gas concentrations (8%, 16%, 24% and 42%). **(b)** Average topographic plots, across all participants (20), of the level-dependent changes in the 15 – 40 Hz spectral scaling exponent (*β*) between an eyes-closed baseline and stepwise increasing end-tidal nitrous oxide gas concentrations (16%, 32% and 47%). Unlike the corresponding level-dependent topographic changes in alpha band power, multiple locations elicit statistically significant group level changes in the spectral scaling exponent (indicated by red asterisks) as determined via a, false discovery rate corrected, nonparametric permutation test (see Methods for further details).

### Experimental level-dependent spectral changes can be accounted for by a superposition of alpha band relaxation oscillatory processes

In the previous section, the spectral changes associated with the administration of gaseous anaesthetic agents (xenon and nitrous oxide) were characterised using alpha band (8-13 Hz) power and the slope (*β*) of the power spectrum (15-40 Hz). Similar changes were induced by both xenon and nitrous oxide in response to stepped increases in gas concentration, namely that topographically significant level-dependent reductions in the spectral scaling, but not alpha band power, occurred. We hypothesise that such changes can be accounted for solely in terms of variations in the underlying distribution of decay rates of a population of, stochastically driven, independent alpha band relaxation oscillators. Such a distribution of decay rates is found by solving an integral equation (see Methods) that describes the experimental power spectrum, over the range 8 – 40 Hz, in terms of the weighted sum of the power spectra of exponentially decaying alpha frequency cosinusoids having a continuous distribution of decay (damping) rates. By using these numerically estimated distribution of dampings to generate model spectra (a forward problem, see Methods for further details), we can assess how well our theoretically motivated decomposition is able to account for the empirically observed, level dependent, spectral changes described in the previous section.

In general we find that these derived model spectra readily reproduce the general spectral features of both baseline, eyes-closed, electroencephalographic activity and the level dependent activity seen in response to xenon and nitrous oxide inhalation. The level-dependent changes in model alpha band power for both xenon (Fig 6a) and nitrous oxide (Fig 7a) replicate well those changes observed in the corresponding experimental spectra. Such a result is also seen with the level dependent changes in the model spectral scaling exponents (Figs 6b & 7b) with the obvious exception being that model spectral scaling exponents are, by construction, always *>* 0.

**Fig 6.**
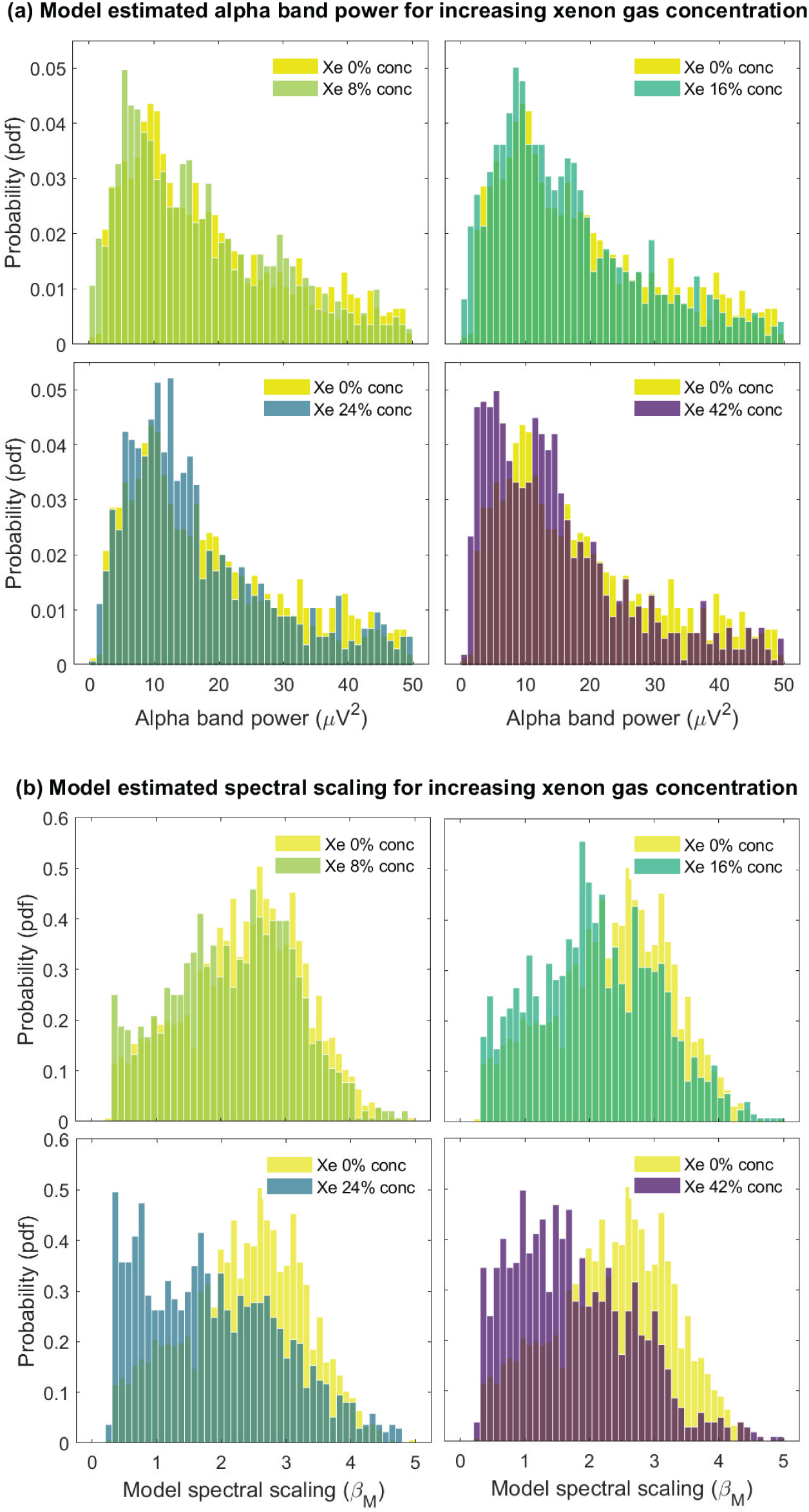
Model estimated distributions of alpha band power and the spectral scaling exponent (*β*) with increasing levels of xenon. (**a**) Normalised histogram of model alpha band (8 – 13 Hz) power, across all electrodes (64) and participants (24), for the eyes-closed baseline and increasing end-tidal concentrations of xenon gas (0% (baseline), 8%, 16%, 24% and 42%). **(b)** Normalised histogram of model 15 – 40 Hz spectral scaling exponents, across all electrodes and participants, for increasing levels of xenon gas (0% (baseline), 8%, 16%, 24% and 42%). These model distributions of alpha band power and spectral scaling exponent (*β >* 0) are consistent with their experimental counterparts (Fig. 1).

**Fig 7.**
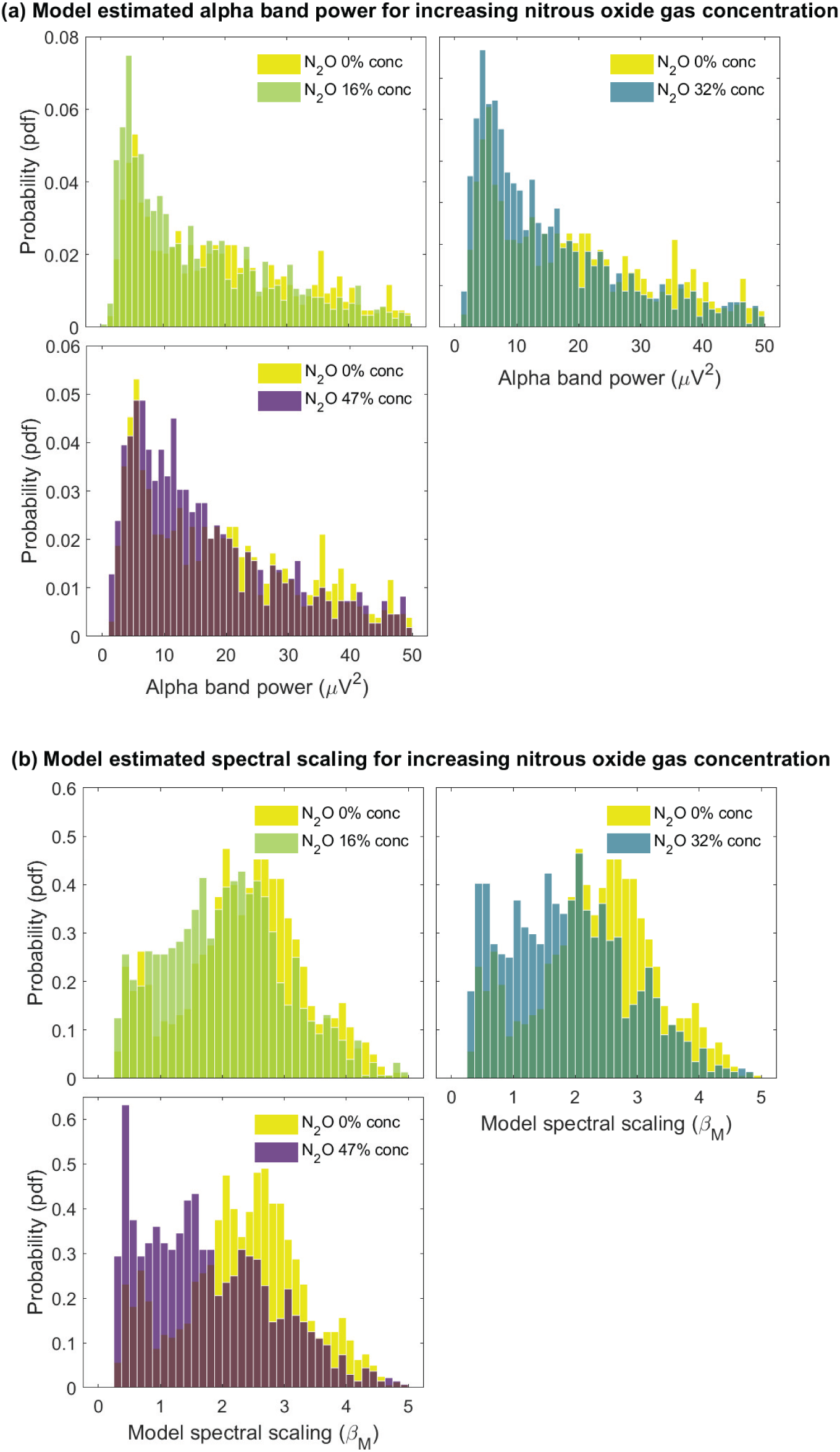
Model estimated distributions of alpha band power and the spectral scaling exponent (*β*) with increasing levels of nitrous oxide. (**a**) Normalised histogram of model alpha band (8 – 13 Hz) power, across all electrodes (64) and participants (20), for the eyes-closed baseline and increasing end-tidal concentrations of nitrous oxide gas (0% (baseline), 16%, 32% and 47%). **(b)** Normalised histogram of model 15 – 40 Hz spectral scaling exponents, across all electrodes and participants, for increasing levels of nitrous oxide gas (0% (baseline), 16%, 32% and 47%). The model distributions of alpha band power and spectral scaling exponent (*β >* 0) for nitrous oxide are consistent with their experimental counterparts (Fig. 2).

Further, scatter plots for experiment versus model alpha band power and slope exponent, across all channels and participants (Fig 8), are strongly correlated. Excluding obvious outliers and the restriction imposed by model scaling exponents having to be greater than zero, the relationship between model and experimental spectral measures is essentially linear.

**Fig 8.**
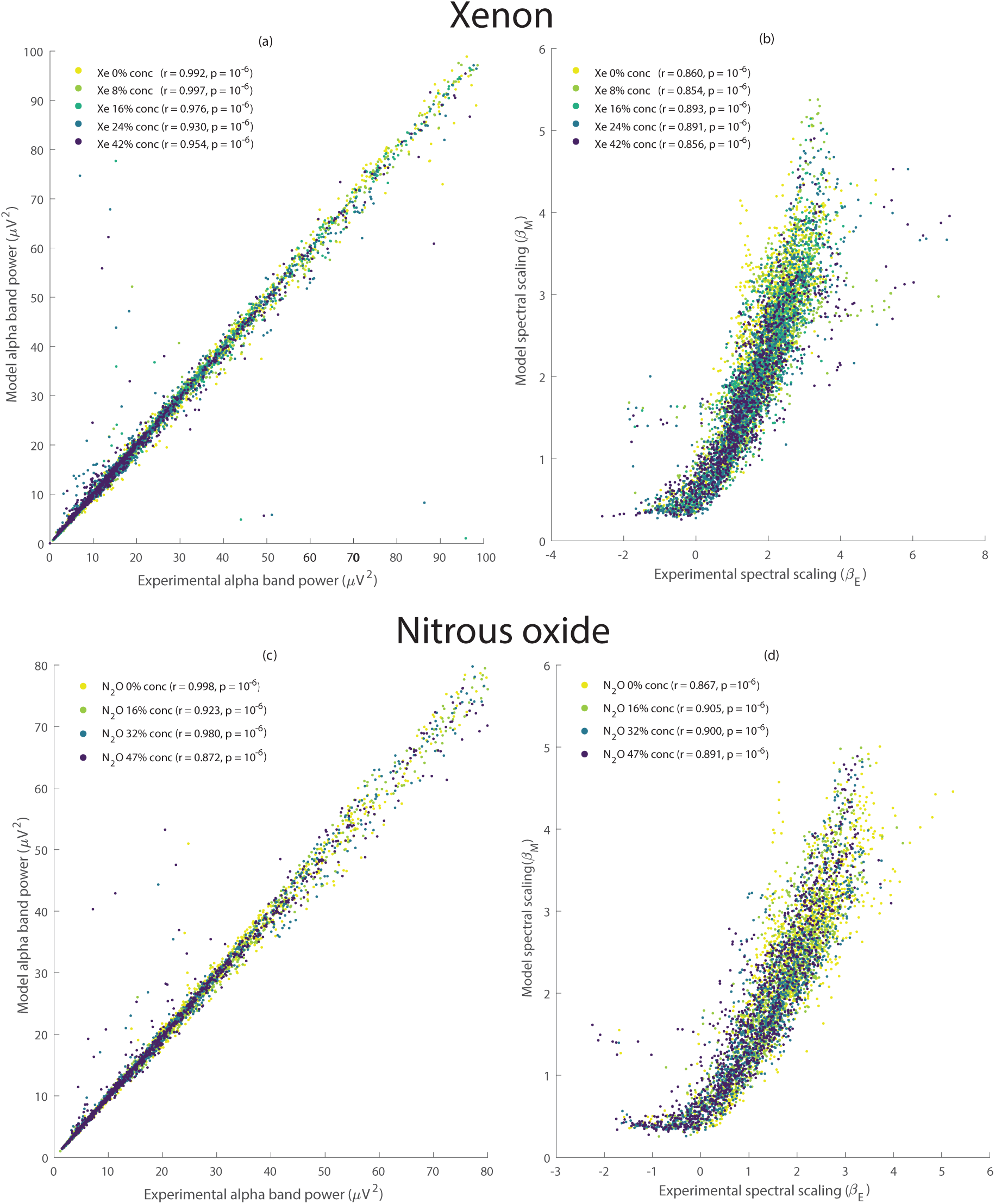
Comparison between model estimated and experimentally recorded spectral features during xenon and nitrous oxide administration. (**a**) Experiment versus model alpha band (8 – 13 Hz) power across baseline and all xenon levels (0% (baseline), 8%, 16%, 24% and 42%). **(b)** As for (a) except for power spectral slope exponent, *β*. **(c)** Experimental alpha band power versus model alpha band power in baseline and across all nitrous oxide levels (0% (baseline), 16%, 32% and 47%). As for (c) except for power spectral slope exponent, *β*. Excluding obvious outliers, there are strong statistically significant (*p* = 10*^−^*^6^) correlations between model and experiment at the population level for both alpha band power and spectral slope exponent. Major deviations from linearity and association between model and experimental scaling are only present for negative spectral slope values. Statistical significance evaluated using non-parametric permutation tests (see Methods for details).

The model decomposition is also capable of capturing the level-dependent topographic changes in alpha band power (Fig 9) and spectral scaling exponent (Fig 10), in particular the graded parieto-occipital changes.

**Fig 9.**
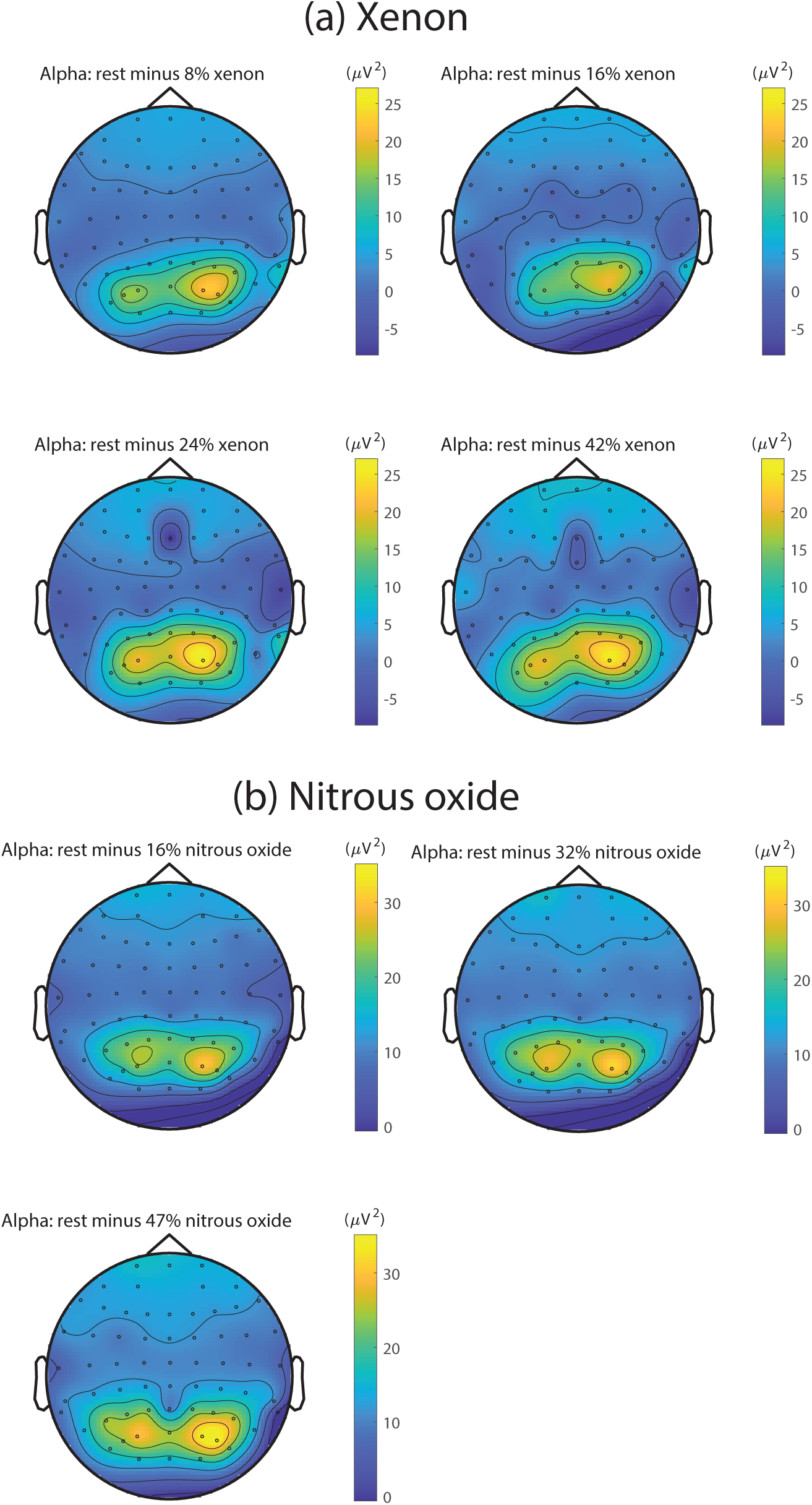
Model estimated topographic variation of group-level alpha band power. (**a**) Topographic maps of group-level differences in model alpha band power between baseline and increasing levels of end-tidal xenon gas levels (8%, 16%, 24% and 42%). **(b)** Topographic maps of group-level differences in model alpha band power between baseline and increasing levels of end-tidal nitrous oxide (16%, 32% and 47%). Model topographic distributions recovers all key experimental features (see Fig 4).

**Fig 10.**
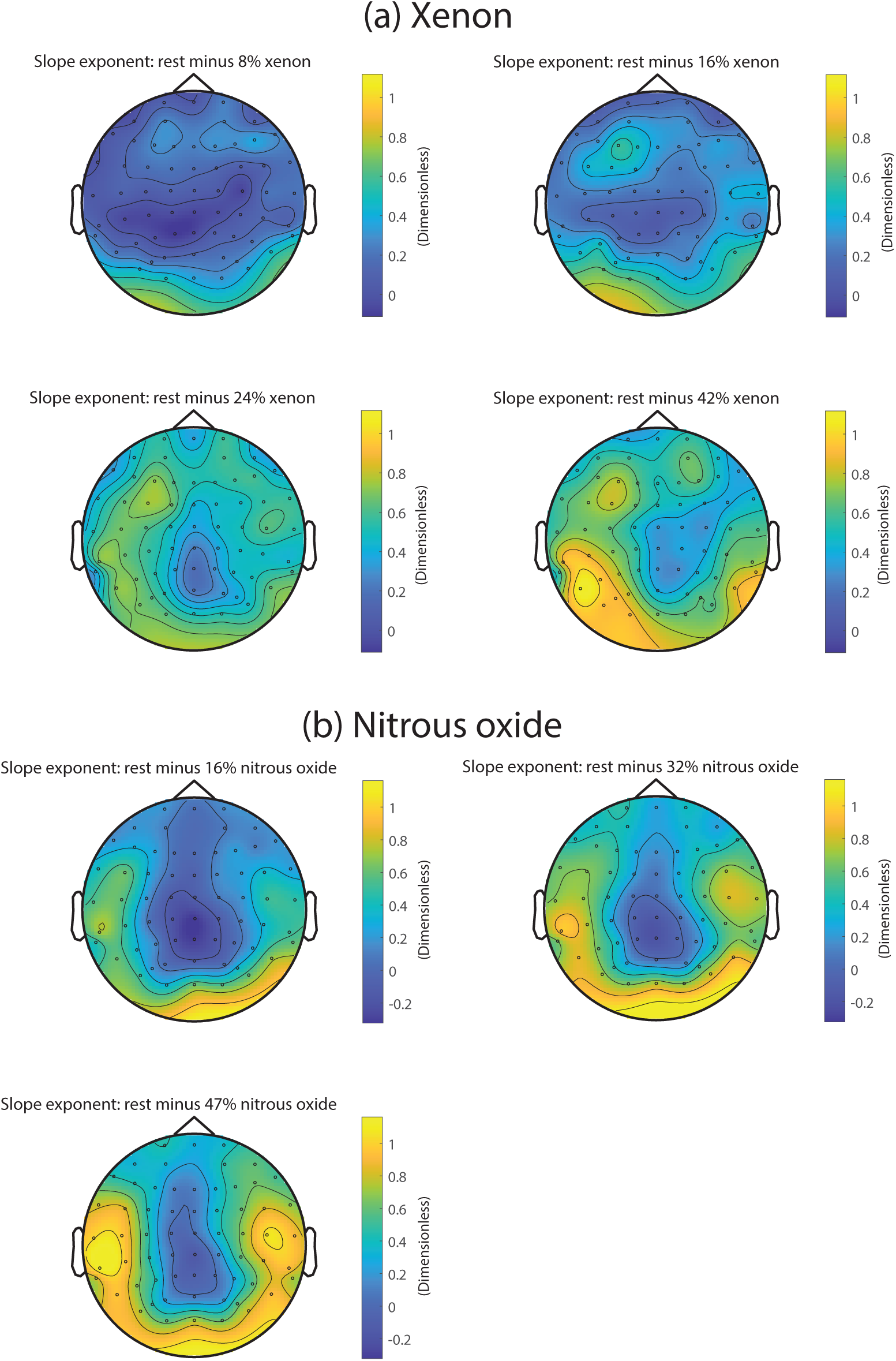
Model estimated topographic variation of group-level spectral scaling. (**a**) **Topographic maps of** group-level differences in model spectral scaling exponent between baseline and increasing levels of end-tidal xenon gas levels (8%, 16%, 24% and 42%). **(b)** Topographic maps of group-level differences in model spectral scaling exponent between baseline and increasing levels of end-tidal nitrous oxide (16%, 32% and 47%). Model topographic distributions recovers all key experimental features (see Fig 5).

As a final demonstration of the fidelity of our theoretical decomposition we investigated the shape of the grand average power spectral density across xenon and nitrous oxide and observed whether it would be reproduced by the corresponding model solutions. The grand average was computed by averaging spectra across all channels and participants for a given agent and level. The corresponding model grand average power spectra were calculated in the same manner by averaging spectra constructed from the numerically estimated damping distributions (see Methods for further details).

For xenon and nitrous oxide we note very similar, level-dependent, broadening of alpha band activity and reductions in peak alpha power (Fig 11 – left column), changes that are clearly observed in the corresponding model spectra (Fig 11 – middle column). The right column of Fig 11 shows the grand average of the numerically estimated damping distributions. Clear level-dependent changes, similar for both xenon and nitrous oxide, are observed. With increasing gas concentration there is an increase in the mean damping and a reduction in broadband cortical driving (Fig 12), as determined by integrating the un-normalised damping distribution *g*(*γ*) (see Eqs. 2-3 in Methods). Broadband cortical driving consistently decreases as end-tidal gas concentration increases (xenon population median: 0.021 [0%], 0.015 [8%], 0.012 [16%], 0.007 [24%] and 0.007 [42%] *µV* ^2^ – nitrous oxide population median: 0.019 [0%], 0.012 [16%], 0.009 [32%] and 0.006 [47%] *µV* ^2^). Pair-wise comparisons show consistent statistically significant (*p <* 0.05) differences at all levels across both xenon and nitrous oxide (Table 4).

**Fig 11.**
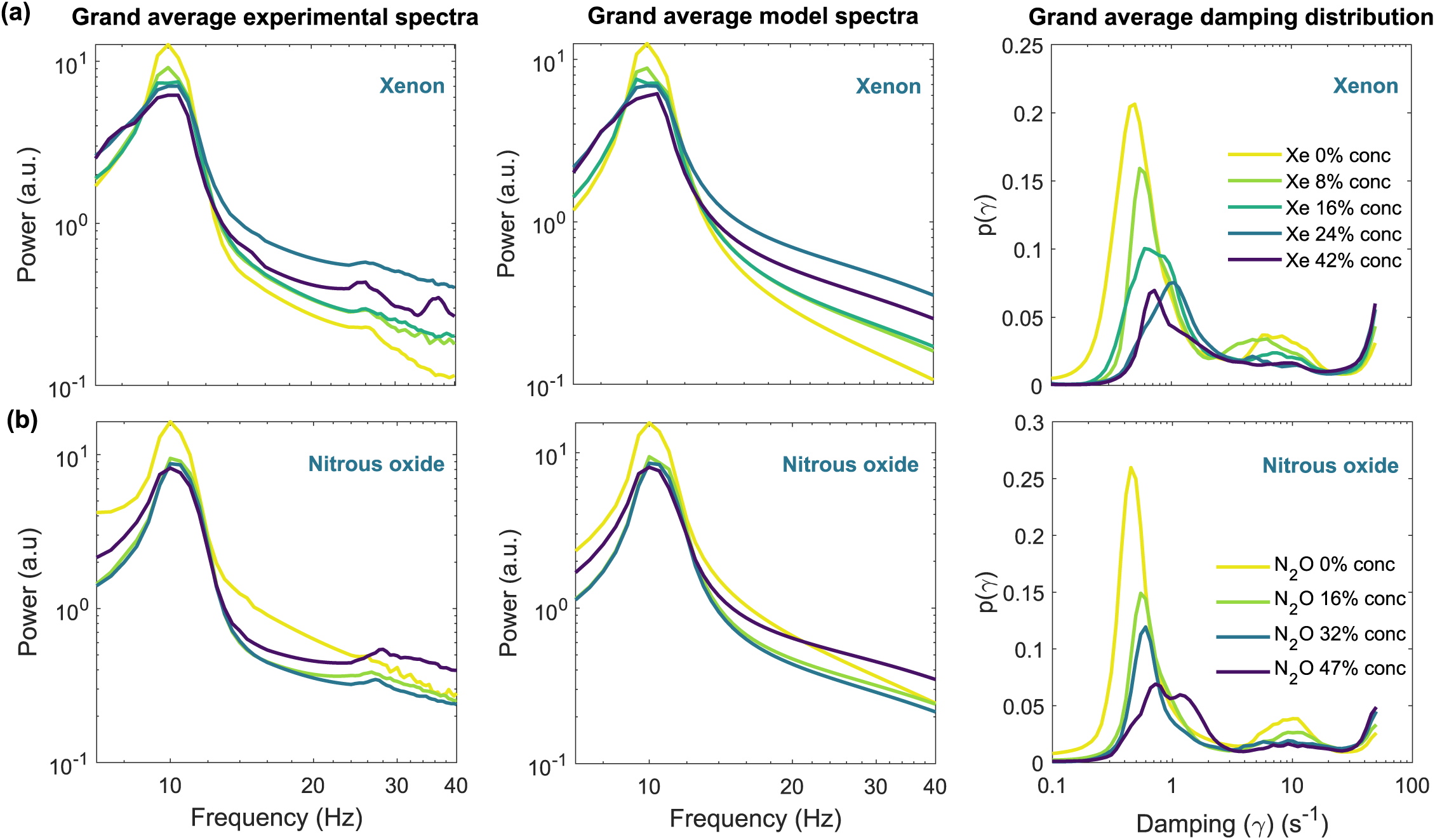
Grand average experimental and model power spectra and the corresponding grand average damping distributions. (**a**) Grand average experimental and model power spectra and damping distributions for baseline and increasing levels of end-tidal xenon gas (0% (baseline), 8%, 16%, 24% and 42%). **(b)** Grand average experimental and model power spectra and the damping distributions for baseline and increasing levels of end-tidal nitrous oxide gas (0% (baseline), 16%, 32%, 47%). The grand averages were taken over all channels and participants.

**Fig 12.**
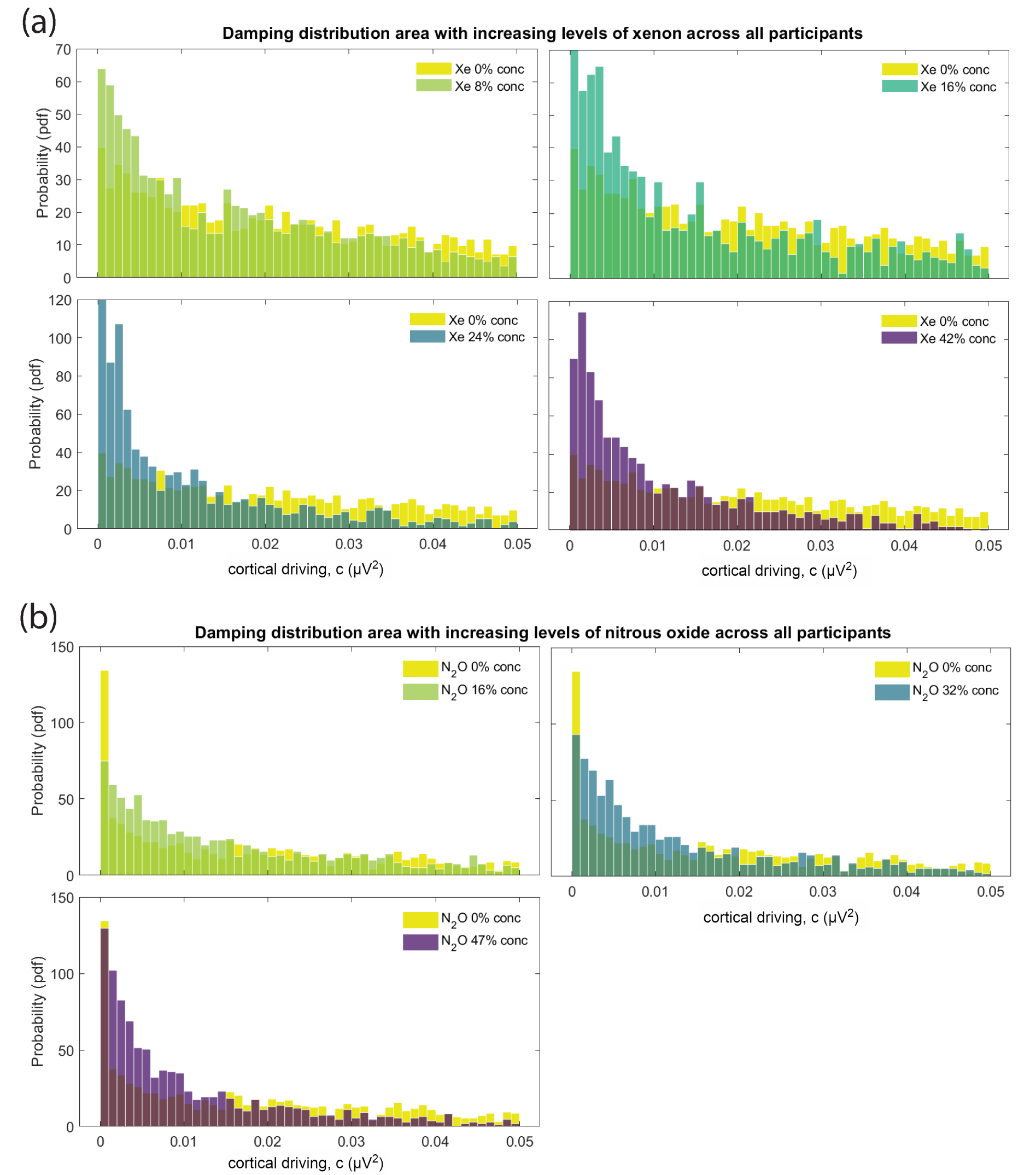
Distribution of broadband cortical driving increasing gas concentration. (**a**) Normalised histograms of the broadband cortical driving (the normalisation constant *c* as obtained by integrating the unnormalized damping distribution *g*(*γ*)), across all participants and channels, for baseline and increasing levels of steady state xenon gas concentration (0% (baseline), 8%, 16%, 24% and 42%). **(b)** Normalised histograms of the broadband cortical driving, across all participants and channels, for baseline and increasing levels of steady state nitrous oxide gas concentration (0% (baseline), 16%, 32% and 47%). When compared to baseline resting state, the distribution of the normalisation constant tends towards smaller values, indicating that the area under the curve increases under anaesthesia, a result observed across both anaesthetic agents.

**Table 4.** Pairwise differences in the median broadband cortical driving for increasing levels of xenon and nitrous oxide. (**a**) Pairwise differences (column – row) in median broadband cortical driving (see Methods) between all xenon conditions (0% (baseline), 8% 16%, 24% and 42%). **(b)** As for (a) except for increasing levels of nitrous oxide (0% (baseline), 16% 32% and 47%). Bolded numbers indicate pair-wise statistical significance (*p <* 0.05) corrected for multiple comparisons (see section Methods for details).

| **(a)** Differences in median broadband cortical driving (xenon) $\mu V^2$ | | | | | |
| --- | --- | --- | --- | --- | --- |
| gas conc | 0 (%) | 8 (%) | 16 (%) | 24 (%) | 42 (%) |
| 0 (%) | 0 | **-0.006** | **-0.009** | **-0.015** | **-0.014** |
| 8 (%) | - | 0 | -0.004 | **-0.009** | **-0.008** |
| 16 (%) | - | - | 0 | **-0.005** | **-0.005** |
| 24 (%) | - | - | - | 0 | -0.001 |
| 42 (%) | - | - | - | - | 0 |
| **(b)** Differences in median broadband cortical driving (nitrous oxide) $\mu V^2$ | | | | | |
| gas conc | 0 (%) | 16 (%) | 32 (%) | 47 (%) |  |
| 0 (%) | 0 | **-0.007** | **-0.010** | **-0.012** |  |
| 16 (%) | - | 0 | **-0.030** | **-0.006** |  |
| 32 (%) | - | - | 0 | **-0.002** |  |
| 47 (%) | - | - | - | 0 |  |

## Discussion

The aims of this research were to determine whether empirically identified spectral alterations of the EEG with increasing concentrations of xenon and nitrous oxide could be accounted for in terms of systematic changes in the distribution of decay rates (dampings) of populations of stochastically driven alpha band relaxation oscillators, as has been previously demonstrated for resting eyes-open and eyes-closed EEG activity [18]. Empirically it was found that increasing levels of both xenon and nitrous oxide resulted in clear dose dependent decreases in alpha band power, peak alpha frequency and the 1*/f^β^*scaling exponent of 15 – 40 Hz activity. Such changes were found to be well modelled by a population of stochastically forced, broad-band driven alpha band relaxation oscillators having a continuous distribution of relaxation rates. Specifically it was found that increasing levels of either gas, both ostensible NMDA receptor antagonists, decreased broadband cortical driving, and the peak alpha frequency, while increasing median alpha band oscillatory damping.

The results of our spectrally based method of estimating alpha band oscillatory damping mirrors and complements results obtained by estimating alpha band damping from raw time series by fitting fixed order ARMA models [22, 23]. Muthukumaraswamy & Liley [23] found that ketamine, another ostensible NMDA antagonist, induced an increase in MEG source level alpha band oscillatory damping that occurred in the context of a flattening of 5-100 Hz power spectral scaling. Liley et al [22] found that increasing levels of end-tidal nitrous oxide during sevoflurane anaesthesia were associated with significant reductions in frontal broadband cortical driving (cortical input), with 66% end-tidal nitrous oxide reducing median cortical input by a factor of approximately 0.5 compared to 100% oxygen. In contrast we found that 47% nitrous oxide reduced the median magnitude of the broadband cortical driving by factor of approximately 0.3, reasonably commensurate when it is noted that we averaged over all scalp electrodes, whereas they recorded from a single frontal electrode in the presence of a potent volatile GABAergic anaesthetic agent.

### Changes in damping, not synchrony, may better explain changes in alpha power

Because of the widespread, and quite reasonable, belief that brain dynamics can only be explicable in terms of the behaviour of individual neurons and their interactions, the idea that EEG amplitude changes are principally driven by alterations in interneuronal synchrony has understandably prevailed. Our results however provide further support for the emerging view that changes in EEG power might be better be explained by alterations in macroscopic neuronal population damping and broadband stochastic driving, rather than by variations in neuronal synchrony. Nevertheless, despite clearly demonstrating that changes in damping and broadband driving can account for the observed changes in EEG power we must exercise some restraint in rejecting the role population neuronal synchrony may have in modulating electroencephalographic power. While it appears, at least in physiologically unperturbed healthy participants [19], that changes in alpha band power between eyes-closed and eyes-open conditions are not associated with any changes in inferred cortical broadband driving, this is not the case when such changes in power are driven by the administration of xenon or nitrous oxide. Because macroscopic decreases in broadband cortical driving and microscopic decreases in interneuronal synchrony are predicted to have the same effect on the shape of the power spectral density [19], we cannot unambiguously determine the role that either of these process have in accounting for the observed empirical changes. Nevertheless, there does appear to exist complementary empirical evidence that favours alterations in broadband cortical input. For example neuroimaging evidence fairly unequivocally demonstrates that xenon and nitrous oxide induce changes in subcortical activity, particularly that of thalamus – a major anatomical source for broadband driving. Xenon has been reported to decrease thalamic regional cerebral blood flow [24, 25] in clinical studies and to disrupt thalamocortical signal propagation, by inhibiting hyperpolarization-activated type 2 cyclic nucleotide-gated channels, in animal models [26, 27]. By contrast nitrous oxide, in the few available studies, is fairly consistently reported to increase thalamic regional cerebral blood flow and glucose metabolism [28, 29]. However unlike xenon nitrous oxide is a potent cerebral vasodilator and thus it is probable that this effect physiologically masks the changes reported for xenon.

### Xenon and nitrous oxide induce reductions in peak alpha frequency and the spectral scaling exponent

The empirical level dependent reductions in alpha band power we identified are broadly consistent with the extant literature. Inhalation of xenon has been reported to produce either negligible effect [1, 2] or notable reductions to alpha band power [4, 6, 30–32]. Similarly, studies involving the administration of nitrous oxide find that resting state levels of alpha band power are preserved [9] or that alpha band power is attenuated [2, 7, 12, 33–35]. Further the reductions we observed in peak alpha frequency have either been directly reported or can be inferred from the redistribution of spectral power. For example McGuigan [36] found frontally recorded peak alpha was reduced by *∼*1 Hz during high levels of inhaled xenon (60%) administered for surgical anaesthesia, a result consistent with Laitio et al [4], who reported that 63% inhaled xenon induced the redistribution of spectral power to slower delta and theta frequency bands such that the spectral edge frequency decreased. Like xenon, nitrous oxide inhalation has been reported to induce a redistribution of spectral power to lower frequencies [7], but as far as we are aware has not been specifically identified as decreasing peak alpha frequency.

In contrast our identified level dependent reductions in the spectral scaling exponent seem to conflict with the few studies that have evaluated it in response to anaesthetic agents [3, 23, 37–41]. Colombo et al [3] evaluated changes in the electroencephalographic scaling exponent in three small (n = 5 each) groups of healthy participants before and during xenon, propofol and ketamine anaesthesia. In the case of xenon anaesthesia, induced by a propofol bolus and maintained at an inspired concentration of approximately 63%, power spectral scaling, calculated over 1 – 20 Hz, 1- 40 Hz and 20 – 40 Hz frequency bands, steepened when compared to an awake eyes-closed baseline, opposite to what we found. However, it can be plausibly speculated that such a steepening represents a methodological aberration given the prior use of a propofol bolus sufficient to induce loss of consciousness and the observation that propofol itself is associated with spectral steepening. Such a conclusion is supported by noting that the other putative NMDA antagonists nitrous oxide [39] and ketamine [3, 23, 38] are reported to reduce spectral slope.

Based on the results of applying our model to empirical data the physiological significance of changes in the spectral scaling exponent is quite different to that proposed by others. In an influential modeling study Gao et al [41] conclude that increases in inhibition, and/or reductions in excitation, causally result in steeper power law spectral scaling i.e., increases in the spectral scaling exponent *β*. In stark contrast we essentially find the opposite. We find that *increases* in median alpha band damping result in shallower (reductions in *β*) power law spectral scaling (see Figs. 6, 7 & 11). In other words we find reductions in *β* reflect a *less* excitable state – rather than a more excitable state when such changes are interpreted in terms of the results of Gao et al [41]. However, similar to a number of other authors [3, 40] they found such steepening in the presence of propofol, a potent GABAergic agent that is well documented to increase alpha band activity [42, 43].

### The observed reductions in peak alpha frequency and the spectral scaling exponent can be theoretically accounted for

Not only does our relationship between scaling and excitability seem more theoretically consistent with the reductions (flattening) in spectral scaling observed during ageing [44–47] and in cognitive impairment [48, 49], the associated changes we observed in peak alpha frequency and model-inferred alpha band damping can be theoretically accounted for by the parametric behaviour of a notable and influential neuronal population model.

Macroscopic neuronal population models [50–52] have emerged as useful alternatives to single neuronal microscopic approaches in exploring the physiological mechanisms underpinning the genesis of electroencephalographic dynamics. Of these models the one of Liley et al [21] is particularly relevant as it purports to account for the mammalian alpha rhythm in the context of a physiologically meaningful parametrisation, such that the phenomenological effects of NMDA receptor antagonism on the emergent alpha rhythm can be investigated. Because the compound excitatory postsynaptic potential (EPSP) consists of both fast *α*-amino-3-hydroxyl-5-methyl-4-isoxazole-propionate (AMPA) and slow NMDA components, NMDA antagonism results in an increase in the EPSP decay rate and a reduction in its peak amplitude [53]. Incorporating these effects into the model of Liley et al [21] results in a reduction in model peak alpha frequency and an increase in alpha band damping [20], effects concordant with what we have reported.

Despite this particular neuronal population model broadly accounting for our results, our results nevertheless challenge the utility of this type of macroscopic theory in characterising resting or time locked EEG activity. The reason is that neuronal population models (also referred to as neural mass or neuronal mean field models) can typically only accommodate a single parametrically homogeneous neuronal population at each spatial location. In contrast our phenomenological approach implies that there are *many* parametrically heterogeneous neuronal populations at each point of the cortical mantle, whose activity is summed to give rise to recorded EEG activity. Thus making parametric inferences, by fitting neuronal population models to real EEG data [54–56], becomes inherently more difficult and uncertain as we *a priori* have no information regarding how many coextensive neuronal populations we should model.

### Global Neuronal Workspace theory: Damping might be related to the likelihood of ‘ignition’

Our results also enable some ‘meta-physiological’ speculation regarding their broader cognitive implications. Since the seminal position-statement of Crick & Koch [57] a plethora of theoretical attempts to account for consciousness have emerged, some of which admit a clear physiological link [58]. Of these arguably the Global Neuronal Workspace (GNW) hypothesis [59] is neurophysiologically the most pertinent. Based on an earlier psychological formulation by Barrs [60], the GNW (also known as the Dehaene–Changeux model) posits that perceptual contents only become conscious when they are widely ‘broadcast’ across the brain by the sudden, coherent and exclusive activation of a subset of neurons, a non-linear process referred to as ‘ignition’. It reasonably follows then, that the less excitable the brain is then the less likely ‘ignition’ will be. Thus, in the context of the GNW theory consciousness is predicted to be impaired if the brain is made less excitable. Indeed this is what our results reveal – increases in levels of the gaseous anaesthetic agents, known to increasingly impair consciousness, reduce whole brain excitability as defined by increases in alpha band damping.

### Limitations of our method

Finally, the inability of our model to incorporate beta band (13 – 30 Hz) activity is an obvious limitation of our approach. While beta band activity exhibits a complex relationship to alpha band activity [61] it is nevertheless clear that they are not independent, and thus reasonably any theoretical method for characterising alpha band activity must reasonably accommodate the characterisation of beta band activity. However, we suspect our simplistic linear approach is not capable of being modified to incorporate beta band activity as we believe, based on a variety of empirical evidence [61–63] that beta band activity does not arise, like alpha, due to the broadband stochastic driving of a stable linear resonance. Nevertheless, it will be important therefore to determine whether any systematic relationship exists between beta band power and alpha band damping, as has been established between alpha and beta band spectral peaks and power spectral densities [61–63].

## Conclusion

In conclusion we have successfully demonstrated that the EEG, in response to the action of xenon and nitrous oxide, can be modelled solely using a distribution of damped alpha band oscillatory processes. While such an approach is inherently simplistic, it is predicated on two well established empirical results regarding the dynamical activity of spontaneous EEG across a range of conditions (rest, sleep, anaesthesia): (i) EEG is not reliably distinguished from a linear random process, and (ii) the EEG has a sharp alpha band resonance/peak, a feature that linearised mean-field models/theories generally account for through a noise driven transfer function having a dominant oscillatory pole within the alpha band. Importantly, the ability of our approach to model the EEG in response to xenon and nitrous oxide provides further empirical evidence that the EEG under anaesthesia, to first approximation, can be considered a filtered pseudorandom linear process, or at the very least be dynamically considered the result of a mixture of populations of damped linear oscillatory processes. Further, this suggests that ‘1/f’ scaling and alpha band oscillatory activity can be dynamically unified, and in doing so, provide an approach that is well equipped to describe the spectral characteristics of the EEG in response to a range of conditions and to provide a foundation for future investigations to build from.

## Methods

The methods follow essentially those of Evertz et al [18]. For clarity, we summarise only the relevant steps, noting any alterations or additions to the original description.

### Modeling EEG as a sum of damped relaxation oscillatory processes

The defining model assumption is that resting state (i.e., spontaneous) EEG originates from the superposition of a large number of uncorrelated, identically stochastically-driven, alpha band damped linear oscillators, which arise from the collective activity of multiple co-extensive cortical neuronal populations. These alpha band processes have a distribution of dampings (relaxation rates), the structure of which is unknown to us. On this basis, our empirically estimated power spectrum, *S*(*f*), is asserted to be decomposable into a sum of power spectral densities, *k*(*f, γ*)), each of which corresponds to the activity of an alpha band relaxation oscillation having a fixed decay (damping) rate, *γ* i.e.,

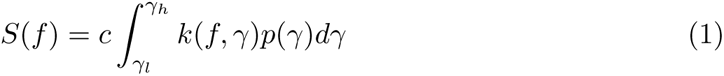

where for simplicity we have chosen to define *k*(*f, γ*) by a Lorentzian-shaped power spectral density function.

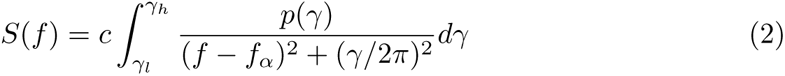

The constant *c* corresponds to broadband cortical driving and *p*(*γ*) defines the distribution of dampings for alpha band oscillations of frequency *f_α_*. Equation 1 corresponds to a Fredholm integral equation of the first kind [64] and thus we seek solutions for *p*(*γ*) given *S*(*f*) and *k*(*f, γ*). Lower (*γ_l_*) and upper (*γ_h_*) integrand limits are selected such that as broad a range of spectral behaviour as possible can be accounted for. For further details see Evertz et al (2022).

### Numerical solution of the model Fredholm Integral equation using non-uniformly weighted Tikhonov regularization

Fredholm integral equations of the first kind are a type of integral equation that are ill-posed, making them highly difficult to solve [65]. To empirically derive useful results, regularization methods are required. We employ the use of Tikhonov regularization (also known as Ridge regression), a well understood method for solving ill-posed problems [65]. Beginning with our model formulation, Eq (2), the Fredholm integral equation of the first kind we choose to solve is

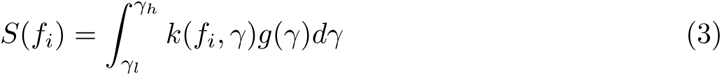

where *g*(*γ*) = *cp*(*γ*). By setting *k_i_*(*γ*) = *k*(*f_i_, γ*), this integral is discretized to give

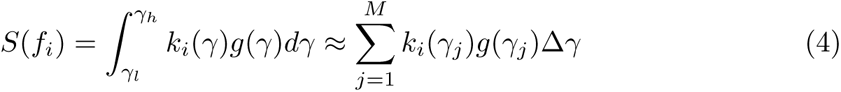

Finally, arranging the above summation as a set of linear equations in matrix notation

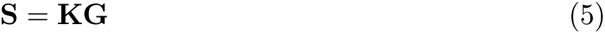

where **S** is an *N ×* 1 data vector (power spectrum), **K** is an *N × M* coefficient matrix with entries consisting of the kernel evaluated at each damping value and **G** is the unknown (un-normalized) distribution vector of dimension *M ×* 1. Simply ‘inverting’ the equation

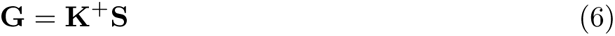

leads to numerically unstable solutions that are sensitive to variations in **S** and those caused by the numerical inversion. However by applying appropriate regularization to Eq (5), we can obtain sensible and numerically stable solutions. Solving the Tikhonov regularization problem involves finding appropriate values for **G** that minimizes the following function for

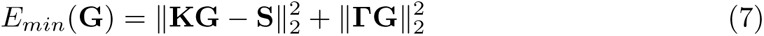

where **Γ** is the *M × M Tikhonov Matrix*, which is generally chosen to be the identity matrix, **I**, multiplied by a regularization parameter, *λ* (i.e. **Γ** = *λ***I** and *|| ·||*_2_ is the Euclidean (or *L*^2^) norm). In practice, Eq (7) is solved as a constrained general least squares minimization problem of the form

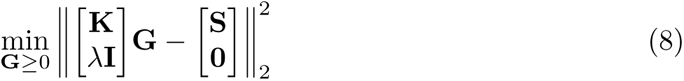

where **G** *≥* 0 means *every* component of **G** is non-negative. We use MATLAB’s lsqnonneg algorithm to solve this optimization problem.

To further enhance the ability to derive high quality spectral fits with this modelling approach, we choose to add a non-uniform weighting when solving Eq (8). The motivation for this is due to how the power is distributed in the EEG. The power contained in the spectrum beyond alpha scales essentially exponentially with frequency, such that lower frequencies will have far larger power than that at higher frequencies. However, when it comes to modelling the power spectral density, minimising the residuals between model and experiment at low frequencies will dominate the optimisation process. Such deviations across residuals that result from data spanning several orders of magnitude is a commonly occurring feature in optimisation problems. This would not present a problem if we are only interested in fitting lower frequencies. However, we wish to also model the high frequency tail as accurately as possible. Therefore, we include a weighting vector to the residuals such that it attempts to weight the low and high frequency residuals evenly. This is achieved by computing a vector of fitting weights and incorporating it into the generalised least squares optimisation routine. The weighting vector is determined by calculating the minimum (*w_min_*) and maximum power (*w_max_*) across all frequency bins of interest for each empirical power spectral density and computing a logarithmically spaced vector of length *N* between the two values i.e.,

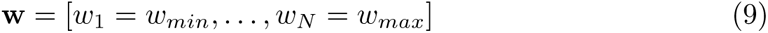

For ease of calculation, we take **w** and diagonalise it into an *N × N* matrix, **W**, with diagonal elements equal to **w** and off-diagonal elements equal to 0. When constructing the general least squares matrix, the weighting matrix is multiplied by both the experimental spectrum input, **S**, which has dimensions of *N ×* 1, and the kernel matrix,

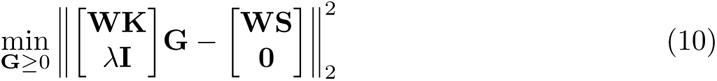

This appropriately weights both the spectrum to be modelled and the kernel matrix which contains the range of profiles to be fitted to the spectrum. The end result of this weighting scheme is that the residuals between the experiment and model, across all frequencies, are more fairly weighted. Thus, when achieving an optimal model fit, reproducing the high frequency tail is now evaluated as being as important as fitting low frequencies around the alpha peak. This optimisation structure was specifically tailored for our purposes, however it was motivated by general weighted optimisations problems described in Aster et al (2018) [65].

### Optimal regularisation parameter selection

Regularisation problems are notoriously challenging to solve and this is compounded by the need to select an ‘optimal’ amount of regularisation. Choosing the appropriate level of regularisation is a careful balancing act between over and underfitting the model. Minimal regularisation can result in poor model fits that are susceptible to the effects of numerical noise from the inversion process, with the opposite true for a high degree of regularisation which can produce results that are over-smoothed. Methods do exist that provide a framework for selecting an optimal regularization parameter. However, these approaches are largely heuristic with ‘optimal’ defined as the value that satisfies the criteria/constraints for the experiment in question [65]. In this study, we explore a regularization parameter range of 10*^−^*^5^ *≤ λ ≤* 10^0^ divided into 100 logarithmically spaced steps, and choose the optimal regularization parameter by finding the maximum entropy solution that falls within an error bound of approximately 2.5% of the minimum residual sum of squares error (RRS) between the model and experimental power spectra. The maximum entropy distribution within that error bound was chosen as it favours solutions that are smoother and broader. See our previous work [18] for a graphical example of the optimal regularization parameter selection process.

### EEG data recorded under administration of nitrous oxide and xenon

Twenty eight healthy male participants were recruited into a study entitled “Effects of inhaled Xe and N_2_O on brain activity recorded using EEG and MEG”, which was approved by the Alfred Hospital and Swinburne University of Technology Ethics Committees (approval No: 260/12). As the experimental procedure follows that described in Pelentritou et al [2], in which the results of a subset of 21 participants were reported, we provide only a brief summary.

Twenty eight healthy male volunteers were administered xenon and nitrous oxide on two separate occasions separated by no greater than 4 weeks. For all sessions, a 5 minute eyes closed resting state baseline was recorded. For both xenon and nitrous oxide, three increasing equi-MAC_awake_ end-tidal gas concentration levels were administered, together with a fourth concentration level for xenon that was designed to induce loss of consciousness in 95% of the participants (xenon: 8%, 16%, 24% and 42%, nitrous oxide: 16%, 32%, 47%). With each increase in gas concentration, a period of approximately 10 minutes was allowed for gas equilibration to occur, before the corresponding 5 minute steady state gas level recording. This procedure was followed for all concentrations across both anaesthetic agents. The EEG was recorded using a 64-channel MEG compatible gel-coupled recording cap at a sampling rate of 512 Hz. We used the EEG portion of the data recorded during the 5 minute baseline eyes closed recording and the 5 minute steady state gas concentration level. Of the 28 participants recruited, 4 of the xenon and 8 of the nitrous oxide recordings were rejected due to excessive artefacts and/or due to the participants inability to tolerate all stepwise increasing gas levels.

### Power spectrum computation and beta band preprocessing

All spectral analysis was performed at the sensor level. Channel-wise power spectral densities for the EEG time series data are calculated by computing Welch periodograms with 2 second Hamming windows with 50% overlap for each relevant 5 minute data epoch. Power spectra activity across a broadband from 7 Hz to 40 Hz was used, as this interval encapsulates both alpha band activity and the high frequency scaling of the spectrum. The choice of the specific upper and lower frequency limits that were used were motivated by consideration of two main factors:

- In resting state EEG the majority (95%) of the spectral power lies below 25 Hz [66]. Therefore, extending the choice of upper limit out to arbitrarily higher frequencies has diminishing returns. By choosing the 40 Hz upper, we also avoid the 50 Hz mains line noise profile.
- Empirical characterisation of the resting EEG has shown the presence of at least two distinct scaling regions; one low frequency and one high frequency with a ‘knee’ frequency generally occurring in the alpha band [3, 23, 67–73]. The specific intervals in each scaling regime vary between studies. However, the consistent feature is that the boundary between these two scaling regions is located within the alpha band and that the two are regarded as arising from dynamically distinct processes. The low frequency portion of the spectrum was excluded as it is considered mechanistically independent of alpha band oscillatory activity.

Due to the individual response of participants to the drugs, there were instances were steady state periods were not available or had excessive artefacts and were rejected from analysis. Therefore, not all participants had recordings for both xenon and nitrous oxide, and not all had recordings at each gas concentration. To characterise the experimental power spectra and possible changes with the levels of anaesthesia, the alpha band power (8-13 Hz), peak alpha frequency and spectral scaling exponent (*β*) (15-40 Hz) were computed across both xenon and nitrous oxide in each condition across all remaining subjects and channels. The alpha band power was calculated by numerically integrating the power spectra across the interval of 8-13 Hz using MATLAB’s trapz. The spectral slope was calculated by fitting a 1*/f^β^* profile to the power spectrum over a frequency interval from 15-40 Hz. Finally, the peak alpha frequency, *f_α_*, was estimated by fitting a single function of the form 1*/*[(*f − f_α_*)^2^ + *b*^2^] to power spectrum alpha band using MATLAB’s curvefit toolbox functions.

Because significant beta band activity (13-30 Hz) typically manifests in the power spectral density as distinct peak/s it may hinder the ability of the model to accurately reproduce the spectral scaling of the high frequency tail. To address this we devised a simple filtering algorithm that removes beta band peaks, yet maintains the spectral scaling of the high frequency tail. This approach is somewhat analogous to the FOOOF/specparam method of [16], where the oscillatory peaks are numerically distinguished from the a 1*/f^β^* arrhythmic background, with ours being simpler in that we were not interested in quantifying beta band activity, but only in removing it and maintaining the scaling of the power spectrum across the region the peak was removed from.

The beta removal routine works as follows: a lower and upper beta band boundary interval is supplied. In each of these boundary intervals (lower: 14-18 Hz, upper: 24-26 Hz) the frequency where the minimum value of the power spectral density occurs is selected. These two frequencies and their associated spectral power values, which are located around the start and end of the classical beta band interval, are used as a fitting interval to which a simple *a/f^β^* profile is fit. From the fitted *a/f^β^* function, a power spectral profile is generated which covers a frequency range between the two selected bounds. The last step involves removing the beta band profile between the two boundaries and replacing it with the generated spectral profile. This removes the beta peak but maintains the approximate spectral scaling. Figure 13 presents a graphical example of the application of the beta removal algorithm. We see that it achieves the desired purpose in the presence of significant beta band activity. Furthermore, it was constructed in such a way that it could be applied systematically to all power spectral densities, regardless of the magnitude of the beta peak, whilst still preserving the appropriate spectral scaling in cases where there is no discernible beta peak.

**Fig 13.**
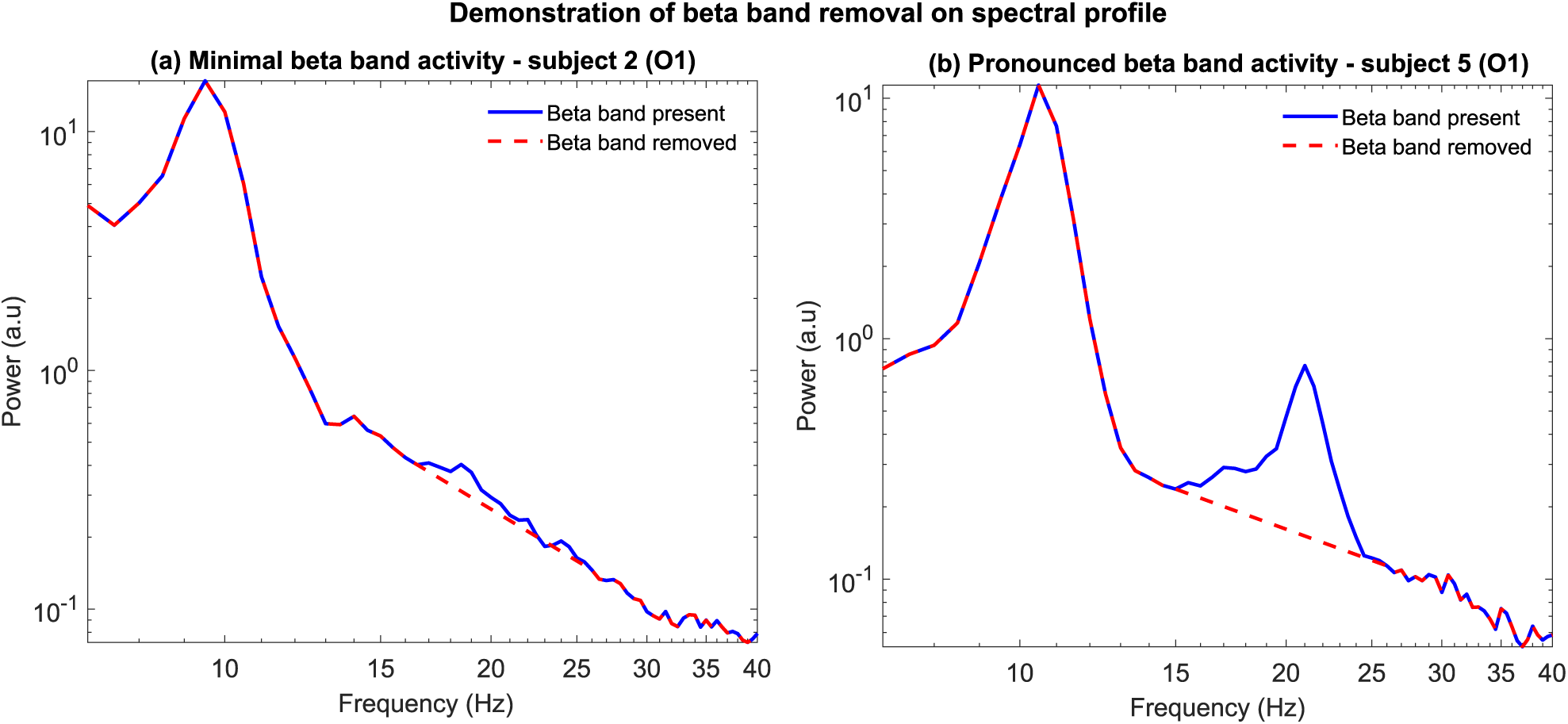
Beta band removal algorithm demonstration. (**a**) Application of beta band peak removal algorithm for a power spectrum with minimal beta band activity (subject 2 – electrode O1). The original spectrum (blue) is plotted alongside the updated spectral profile (dashed red). We note that when minimal beta band activity is present, the algorithm has little effect on the power spectrum. **(b)** Same as (a) except in this case the participant (subject 5 – electrode O1) has a large beta band peak which is effectively removed with application of the algorithm.

### Statistical analysis

Significant differences between the relevant level-dependent empirical and model distributions were determined by 1) Estimating the relevant distribution median values and the difference between conditions 2) Perform a Kruskal-Wallis (one-way ANOVA by ranks) test to determine whether any significant level-dependent changes and 3) Performing multiple pair-wise comparisons between each relevant pair of gas conditions and conservatively correcting for the multiple comparisons using a Bonferroni correction, such that only robust differences are identified.

